# Growth patterns of caudal fin rays are informed by both external signals from the regenerating organ and remembered identity autonomous to the local tissue

**DOI:** 10.1101/2024.03.29.586899

**Authors:** Melody Autumn, Yinan Hu, Jenny Zeng, Sarah K. McMenamin

**Affiliations:** Biology Department, Boston College, Chestnut Hill, MA 02467

**Keywords:** Regeneration, Zebrafish, Transplant, Growth rate, Skeletal patterning, Thyroid hormone

## Abstract

Regenerating tissues must remember or interpret their spatial position, using this information to restore original size and patterning. The external skeleton of the zebrafish caudal fin is composed of 18 rays; after any portion of the fin is amputated, position-dependent regenerative growth restores each ray to its original length. We tested for transcriptional differences during regeneration of proximal versus distal tissues and identified 489 genes that differed in proximodistal expression. Thyroid hormone directs multiple aspects of ray patterning along the proximodistal axis, and we identified 364 transcripts showing a proximodistal expression pattern that was dependent on thyroid hormone context. To test what aspects of ray positional identity are directed by extrinsic cues versus remembered identity autonomous to the tissue itself, we transplanted distal portions of rays to proximal environments and evaluated regeneration within the new location. While neighboring proximal tissue showed robust expression of *scpp7*, a transcript with thyroid-regulated proximal enrichment, regenerating rays originating from transplanted distal tissue showed reduced (distal-like) expression during outgrowth. These distal-to-proximal transplants regenerated far beyond the length of the graft itself, indicating that cues from the proximal environment promoted additional growth. Nonetheless, these transplants initially regenerated at a much slower rate compared to controls, suggesting memory of distal identity was retained by the transplanted tissue. This early growth retardation caused rays that originated from transplants to become noticeably shorter than their native neighboring rays. While several aspects of fin ray morphology (bifurcation, segment length) were found to be determined by the environment, regeneration speed and ray length are remembered autonomously by tissues, persisting across multiple rounds of amputation and regeneration.

**GRAPHICAL ABSTRACT:** 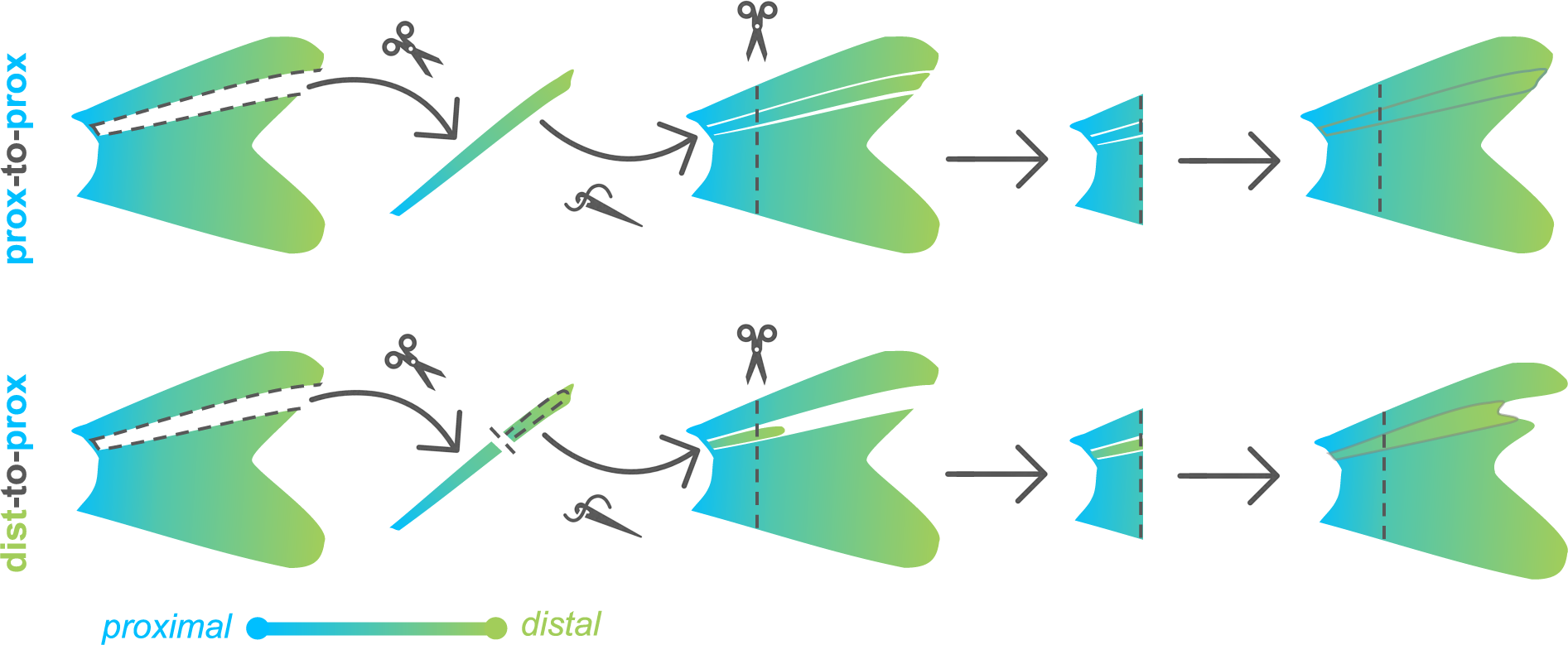

## INTRODUCTION

To restore the original morphology of an appendage, regeneration must faithfully rebuild lost tissue. The morphology and size to which regenerating tissue grows must be determined by positional information (Wolpert, 1969). Such cues could be informed by remembered positional identity or could be interpreted from environmental cues from surrounding tissue (e.g. diffusible or spatially distributed factors). However, these two potential inputs can be difficult to disentangle.

Zebrafish fins are powerful models for studying regeneration and can provide new insights into the nature of positional memory and the pathways that regulate regional growth and patterning. The caudal fin is made up of symmetrical dorsal and ventral lobes, each composed of nine segmented fin rays. Upon amputation, a blastema of de-differentiated cells forms (Knopf et al., 2011; Tu & Johnson, 2011), and each ray regrows from the wound site to rebuild its original morphology (reviewed in Harris et al., 2021; I. M. Sehring & Weidinger, 2020).

Regeneration rate is informed by the relative proximodistal location of the regenerating tissue on the fin (Lee et al., 2005). Distal amputations are followed by slow regenerative growth, while proximal amputations close to the body initiate rapid growth that progressively slows as the regenerate approaches the original size (Akimenko et al., 1995; Banu et al., 2022; Lee et al., 2005; Uemoto et al., 2020). Regardless of how much tissue is removed, regeneration restores the organ to its original size within three weeks (Wehner et al., 2014).

Intact fin rays exhibit morphological differences along the proximodistal axis. At the proximal base, ray segments are longest and widest, tapering and shortening progressively towards the distal edge; rays also form bifurcations at specific locations along the axis (Harper et al., 2023). Components of proximodistal patterning are regulated by thyroid hormone (TH), which induces distal features (Harper et al., 2023). Proximal and distal tissues from intact adult fins show unique transcriptomic profiles (Rabinowitz et al., 2017), and these expression patterns are regulated by developmental TH (Harper et al., 2023). Here, we tested if transcriptomic differences are apparent during the regeneration of proximal compared to distal regions of the fin.

The relative length of individual rays appears to be remembered autonomously by tissues (Uemoto et al., 2020). Fin rays differ in length from the central to the peripheral regions of the fin, giving the organ an overall forked shape. Previous transplantation experiments demonstrate that when short central rays are swapped with long peripheral rays, the tissue regenerating in the new environment produces a ray of intermediate length (Shibata et al., 2018). However, it remains unclear whether proximodistal location along an intact ray imprints remembered positional information that could inform morphology during regeneration.

Transplants of blastema cells from different proximodistal locations were not able to influence lengths of regenerates (Shibata et al., 2018). Further, hemi-rays—the apposed contralateral bones that make up individual ray segments—can be transplanted to different proximodistal locations; the resulting recombinant rays regenerate with morphologies expected for the regenerating environment (Murciano et al., 2007). Nonetheless, given the notable differences in gene expression and morphology along the intact proximodistal axis, we asked if entire ray segments could remember proximodistal identity, and we tested the ability of this memory to influence gene expression, regrowth rates, ultimate length and patterning of regenerating rays.

## RESULTS

### Regenerating fin tissue shows unique proximodistal transcription

Many genes show proximodistal differences in expression as the caudal fin regenerates (Akimenko et al., 1995; Lee et al., 2005), and we aimed to capture trends across the transcriptome during the regeneration of the organ. Amputating at a consistent proximal location, we evaluated expression from three regions as they regenerated a proximal region collected after the blastema had already formed (Wang et al., 2019; 4 days post amputation, dpa), a middle region midway through regeneration as the ray bifurcations were forming (7 dpa), and a distal region (15 dpa; see Supplementary Fig. 1C). We identified 489 transcripts that were differentially expressed between proximal and distal regenerating tissue (Fig. 1A-B): 29 genes were proximally enriched and 460 were distally enriched. GO term analysis of differentially expressed transcripts along the proximodistal axis showed enrichment of genes involved in pigmentation, likely reflecting differentiation of pigment cells (Supplementary Fig. 1G). Transcripts dependent on thyroid hormone were enriched for gas transport GO terms, potentially reflecting shifts in circulation and metabolism (Supplementary Fig. 1H). In our regenerates, ray bifurcations were actively forming during the middle time point (7 dpa); however, this tissue revealed no transcripts that were differentially expressed compared to proximal and distal tissues.

**Figure 1.**
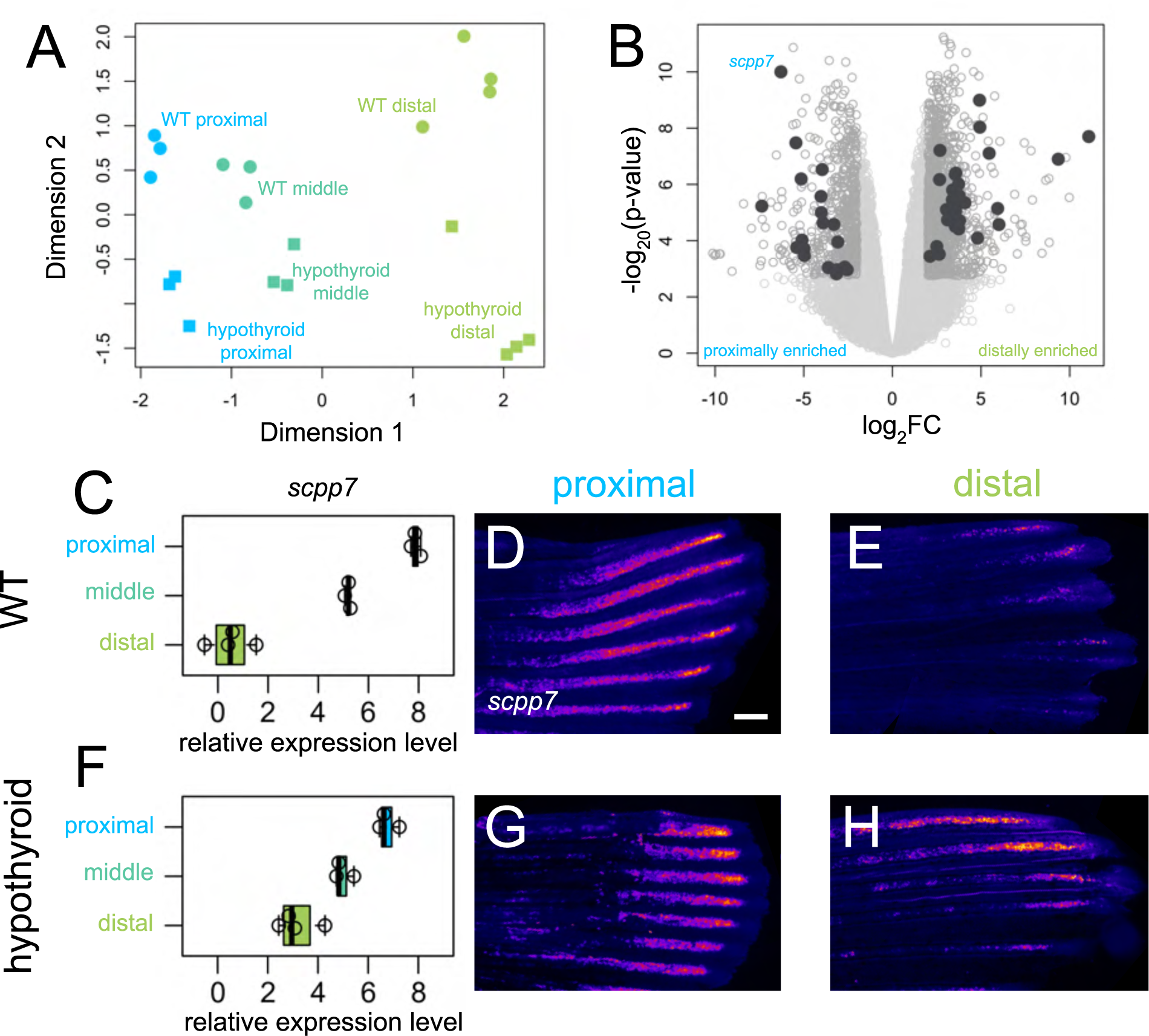
Thyroid hormone distalizes gene expression patterns during regeneration. (A) Multidimensional scaling plot comparing gene expression profiles in different regions (proximal, 4 dpa; middle, 7 dpa; distal, 15 dpa) of regenerating tissue from WT and hypothyroid fish; each data point represents one biological replicate. (B) Volcano plot showing differential gene expression between regenerating proximal and distal regions in WT. Filled grey circles indicate thyroid hormone-dependent genes. (D, F) *scpp7* relative expression in (C) WT and (F) hypothyroid tissue samples. Note increased proximal expression in hypothyroid distal tissues. Whole mount fluorescent *in situ* hybridization using custom *scpp7* RNAscope probe on (D-E) WT and (G-H) hypothyroid tissue regenerating (D, G) proximal or (E, H) distal fin tissues. Warm colors indicate highest regions of expression. Scale bar, 200 μm

### Hypothyroid tissues lose proximodistal differential expression of many genes

Developmental hypothyroidism proximalizes both transcriptional expression and ray morphology in intact fins (Harper et al., 2023), and we asked if fins regenerating in a hypothyroid context also showed proximalized gene expression patterns. We reasoned that transcripts with a TH-regulated expression differential would be strong candidates as mediators of proximodistal patterning and distal identity. Analyzing the transcriptomes holistically, the major axes of variation robustly captured proximodistal location (dimension 1) and TH condition (dimension 2), but there was little apparent correlation between the two (Fig. 1A). Nonetheless, certain transcripts showed a proximodistal differential in expression that was dependent on the presence of TH. Indeed, of the 489 differentially expressed genes found in WT tissue, 364 lost proximodistal specificity in hypothyroid tissue: ∼86% (25/29) of proximally enriched and ∼76% (349/460) of distally enriched genes lost proximodistal differential expression in a hypothyroid context.

### *scpp7* is proximally enriched during regeneration

Of the transcripts showing TH-dependent proximal enrichment, secretory calcium-binding phosphoprotein 7 (*scpp7*) could be robustly visualized using RNAscope (Fig 1D-E, G-H). Along with other SCPP factors, SCPP7 is involved in bone mineralization (Kawasaki, 2009), and is strongly upregulated during scale regeneration (Bergen et al., 2022). Proximal tissues showed robust expression of *scpp7* in both WT and hypothyroid backgrounds, but the gene was more strongly expressed in distal tissue from hypothyroid regenerates compared to those of WT (Fig. 1C-H).

We asked if the variation in *scpp7* expression in the different regions could be attributed to differences in the time since injury rather than proximodistal position of regeneration. To test this possibility, we performed distal amputations on WT fins (see Supplementary Fig. 1D) and assessed *scpp7* expression in 4 dpa distally-regenerating tissue. *scpp7* expression was similar to that of 15 dpa distally-regenerating tissues (Supplementary Fig. 1E-F), suggesting that this expression differential indeed characterizes distal regenerating tissue.

### *scpp7* expression in regenerating tissues reflects original proximodistal location rather than regenerative environment

We asked whether attenuated *scpp7* expression would be remembered by distal tissues if they regenerated in a proximal context. To test this, we designed a distal-to-proximal ray transplantation procedure in which a ray was removed from the fin, and a distal portion of the extirpated ray was transplanted into the proximal position.

After the distal transplant integrated into the proximal location, the entire fin was amputated (through the transplant) to allow distal tissue to regenerate alongside proximal tissue (“dist-to-prox”, Fig. 2A-C; see Methods and Supplementary Fig. 10 for additional details). A completely extirpated ray with no transplant produced no regeneration (Supplementary Fig. 2). We assessed *scpp7* expression in the regenerate originating from the dist-to-prox transplant and found expression was significantly reduced compared to those of neighboring proximal rays at 4 dpa (Fig. 2D-E). This recapitulation of distal-like expression while regenerating in a proximal context suggests that expression level of this transcript is informed by original tissue identity.

**Figure 2.**
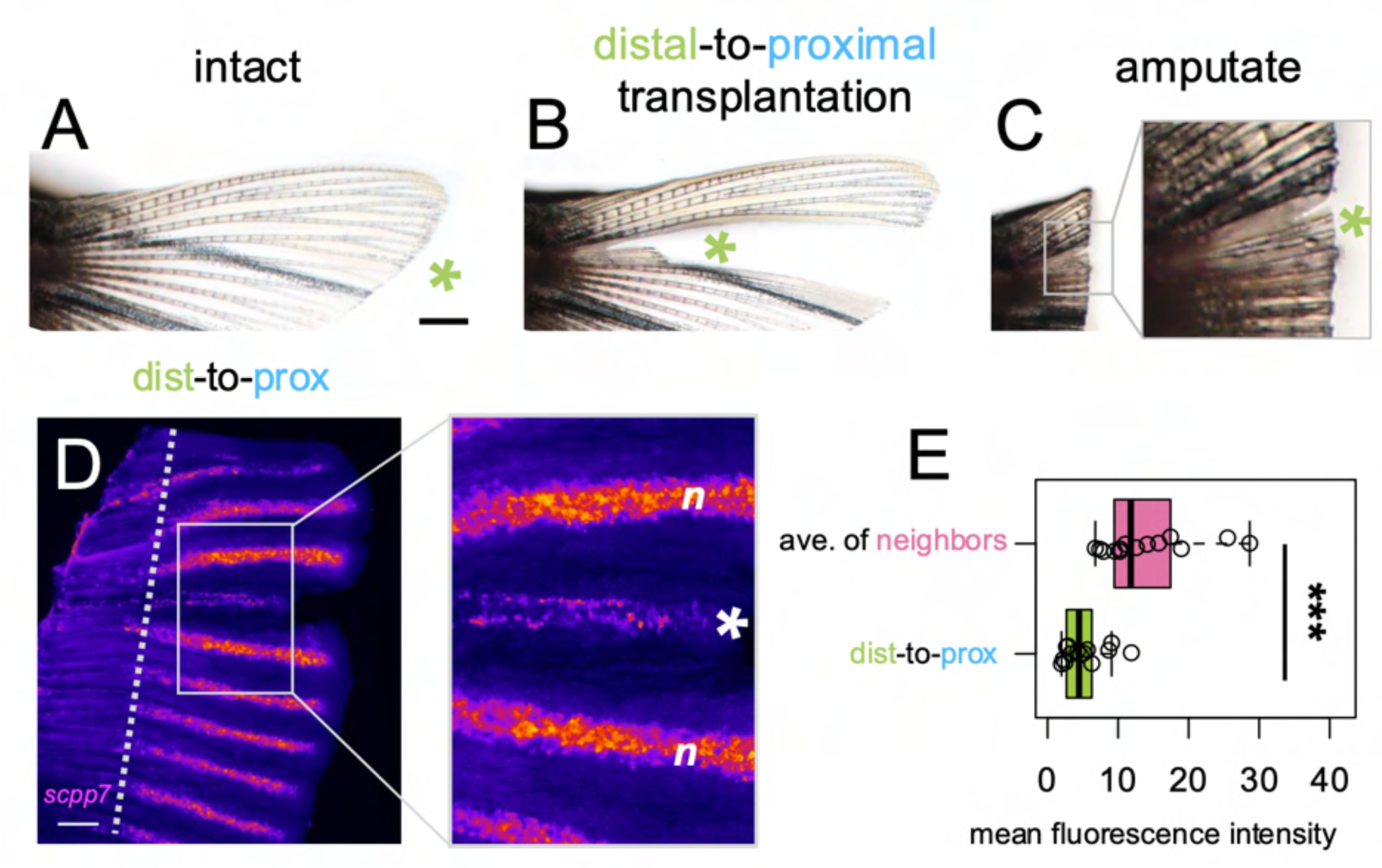
*scpp7* expression in regenerating tissues reflects original position rather than current environment. (A-C) Example of a fin lobe subjected to the distal-to-proximal transplantation procedure. (D) Whole mount fluorescent *in situ* hybridization with *scpp7* RNAscope probe on dist-to-prox regenerating fins at 4 dpa. Warm colors indicate highest regions of expression. (E) Boxplot showing mean fluorescence intensity of dist-to-prox transplant tissue (asterisk) and the average intensity of its peripheral-most and center-most neighbors (n). Significance determined by a Welch two-sample paired t test. Scale bars, (A) 1 mm; (D) 200 μm.

### Distal-to-proximal transplanted tissue restores shorter fin rays

We predicted that if dist-to-prox transplanted tissue possessed remembered positional identity, precocious distal features should be apparent in the resulting regenerate. To adequately evaluate subtle differences in regrowth, we needed a comparison that had undergone identical microsurgery without introducing a major axis translocation. Thus, we performed control “prox-to-prox” transplants, extirpating a ray, then grafting the entire tissue back into its position (Fig. 3A-D). Interestingly, these prox-to-prox rays were not able to regenerate to the same length as the corresponding rays on the ventral lobe (Supplementary Fig. 3K) and were ultimately slightly shorter than undisturbed neighboring rays. During microsurgery, prox-to-prox rays inevitably lost 1-3 segments and about a mm in length (Supplementary Fig. 3I), so the change in ultimate length may reflect a slight positional shift. Additionally (or alternatively), the manipulation of the microsurgery itself may be sufficient to effect patterns of regeneration.

**Figure 3.**
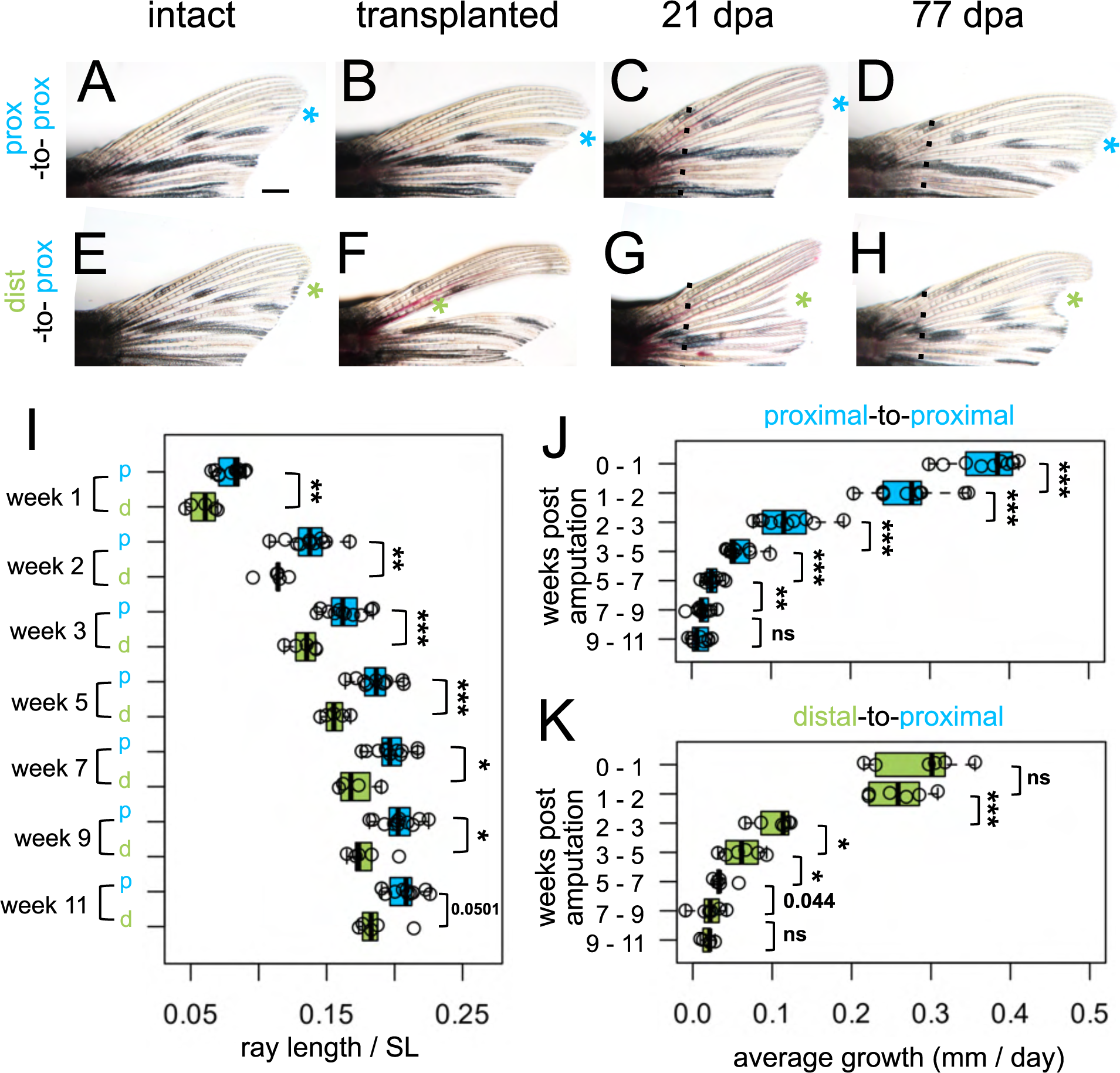
Regrowth rate reflects both intrinsic identity and the regenerative environment. Dorsal fin lobes of (A-D) proximal-to-proximal (blue asterisk) or (E-H) distal-to-proximal (green asterisk) transplantation: (A, E) intact pre-transplantation, (B-F) one day post-transplantation, (C-G) regenerating at 21 dpa, (D, H) regenerating at 77 dpa. Amputation plane, dashed line. (I) Prox-to-prox versus dist-to-prox ray length (measured from amputation plane) normalized by standard length (SL) for each week. Average amount of growth per day during different growth periods for (J) prox-to-prox or (K) dist-to-prox rays. Significance determined by (J-K) paired or (I) unpaired Welch two-sample t tests. Scale bar, 1 mm.

Compared to the microsurgery-controlled baseline of prox-to-prox rays, rays originating from dist-to-prox transplants were consistently shorter, through eleven weeks after amputation (Fig. 3I). Dist-to-prox regenerates were obviously shorter than both neighboring rays (Fig. 3G-H) and the corresponding ray on the ventral lobe (Supplementary Fig. 3G-H). These differences in ultimate length suggest that the dist-to-prox transplants autonomously retain memory of their original proximodistal identity.

### Growth rates during regeneration reflect both intrinsic identity and environmental context

Since the dist-to-prox regenerates were significantly shorter compared to prox-to-prox regenerates, we asked whether these regenerates grew at a relatively slower pace. During the first week of regeneration (weeks 0-1), prox-to-prox transplants regenerated rapidly, adding 2.6 mm (0.37 mm per day); in contrast, dist-to-prox regenerates grew much more slowly, adding only 2.0 mm during this first week (0.27 mm per day; Fig. 3J-K). By the second week (weeks 1-2), the two types of transplants were growing at comparable speeds, adding 1.9 mm length (0.27 mm per day). Through the remainder of the eleven-week period, dist-to-prox and prox-to-prox rays maintained similar regrowth speeds (Fig. 3J-K). Growth rates plateaued after week nine, as the regenerates reached isometric growth (Fig. 3J-K). Prox-to-prox rays’ regrowth speed was reduced in comparison to corresponding ventral rays during the first week of regeneration but by the second week they kept pace (Supplementary Fig. 3J-K).

### Fin ray patterning is environmentally coordinated

Bifurcations are a discrete indicator of proximodistal morphology (Harper et al., 2023). We asked whether the origin of tissue (distal versus proximal) would influence the location of the bifurcation in a regenerate, and we quantified the bifurcation position in dist-to-prox and prox-to-prox rays. Bifurcations formed in the location expected for the environment regardless of transplant type (Fig. 4C), suggesting bifurcation position is the result of globally coordinated cues (Dagenais et al., 2021; Murciano et al., 2002, 2007) rather than being locally regulated by tissues based on remembered identity.

**Figure 4.**
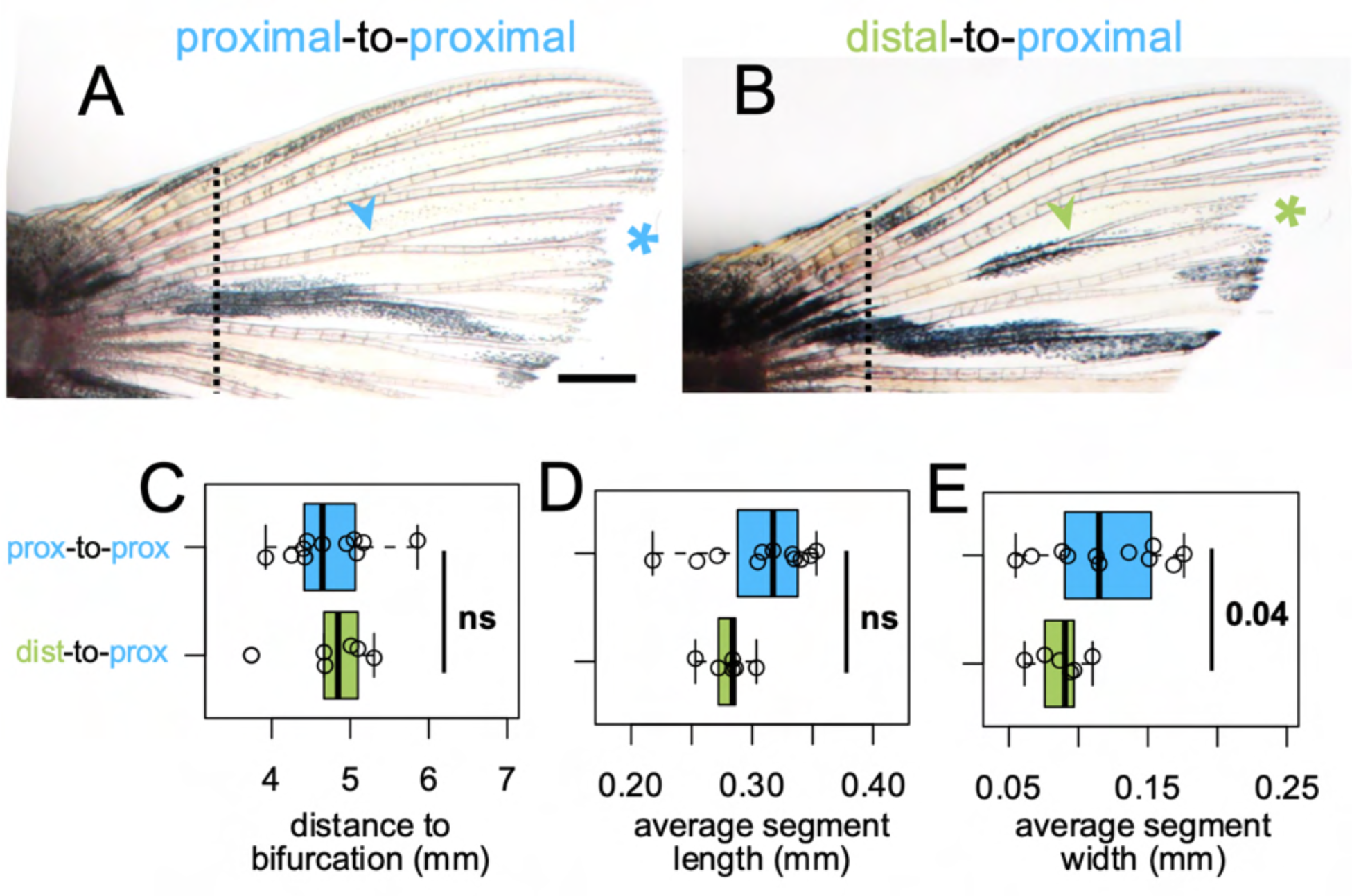
Fin ray patterning matches environment. Dorsal fin lobe at 35dpa after either (A) proximal-to-proximal (blue asterisk) or (B) distal-to-proximal (green asterisk) transplantation. Amputation plane shown with dashed line. Arrowheads indicate primary bifurcations. Boxplots showing the (C) proximodistal position of the bifurcation, (D) average segment length, and (E) average segment width in regenerate. Significance determined by a Welch two-sample t test. Scale bar, 1 mm.

In evaluating appropriate controls for our dist-to-prox transplants, we discovered that while ray length is similar between dorsal and ventral lobes, the proximodistal patterning differs between the dorsal and ventral lobes of uninjured fins (Supplementary Fig. 4). Further, regenerated ray segments were somewhat longer and wider than segments of the intact fin, with bifurcations farther from the body (Supplementary Fig. 5E-F; Supplementary Fig. 6I-J; also see Azevedo et al., 2012). Therefore, we the prox-to-prox transplants were used as the best comparisons for dist-to-prox proximodistal patterning. While dist-to-prox rays regrew marginally thinner segments, segment length was comparable to that of prox-to-prox ray segments (Fig. 4D-E).

The total length of a regenerating fin can be increased by treating with a calcineurin inhibitor, but these treatments do not alter positional memory and rays return to their WT baseline upon regeneration (Daane et al., 2018). We asked if regenerating from a pharmacologically-lengthened segment would alter the patterning of a regenerating ray; however, once calcineurin inhibition was removed, segment length and location of bifurcation were indistinguishable from control regenerates (Supplementary Fig. 8).

### Rays originating from distal transplants remember their length through multiple rounds of regeneration

To test whether the intermediate length of dist-to-prox rays would be remembered, we performed multiple rounds of regeneration, amputating distal to the previous amputation plane (Fig. 5A-D). Even after three rounds of regeneration, rays originating from dist-to-prox transplants were always significantly shorter than corresponding ventral rays (Fig. 5E-G).

**Figure 5.**
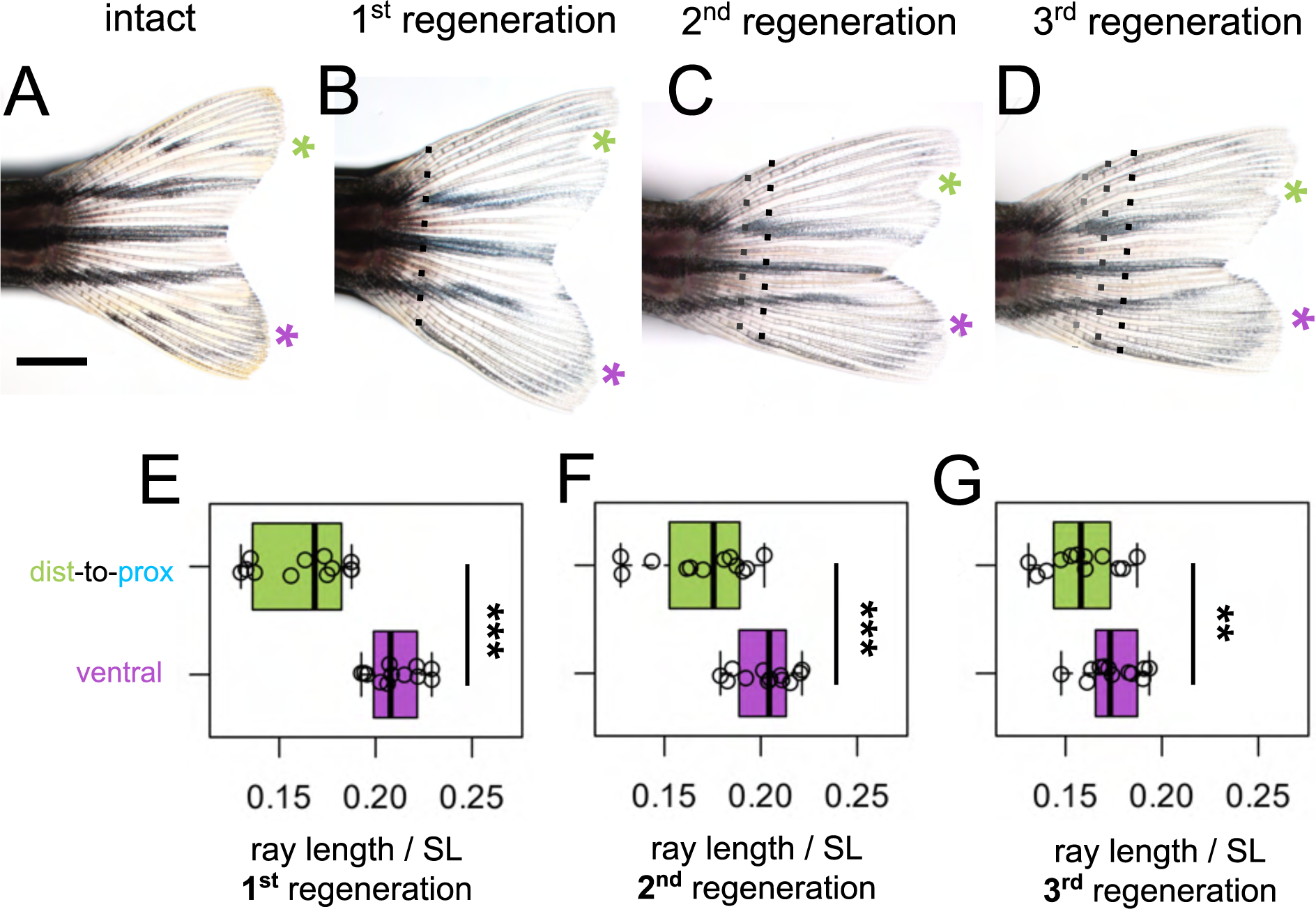
Shorter ray length is remembered through multiple regeneration cycles. (A) Intact fin. (B-D) Regenerating fin after distal-to-proximal transplantation: (B) 28 days post first amputation, (C) 28 days post second amputation, (C) 28 days post third amputation. Green asterisk, dist-to-prox; purple asterisk, ventral ray. Black dashed line, most recent amputation. Grey dashed lines, previous amputation planes. Boxplots showing the total length standardized by SL after (E) first, (F) second, and (G) third regeneration. Ray length was measured from the most recent amputation plane. Significance determined by paired Welch two-sample t test.

## DISCUSSION

Intact fins show transcriptomic differences across the proximodistal axis (Harper et al., 2023; Rabinowitz et al., 2017), and here we identified a suite of genes that shift as the fin regenerates different proximodistal regions. Previous transcriptomic analyses of regenerating fins have focused on the early shifts in expression as the tissue initiates regenerative regrowth (Li et al., 2021; Nauroy et al., 2019); we found that there are substantial shifts in expression patterns even after regeneration is underway, corresponding with different stages of outgrowth proceed (see Akimenko et al., 1995; Lee et al., 2005). Indeed, we found ten times as many distally-as compared to proximally-enriched transcripts; this may reflect the increased number of differentiated cells in the more mature regenerate (Nauroy et al., 2019).

The presence of TH throughout development distalizes gene expression patterns as fins develop (Harper et al., 2023); however, during regeneration, hypothyroidism did not proximalize pattern of gene expression. It is possible that temporal shifts in the regenerating transcriptome overwhelmed any TH-dependent proximodistal pathways in our analysis, since the three stages analyzed vary in both time since injury and proximodistal regions of regeneration. Nevertheless, we identified many genes expressed in a proximodistal differential that was dependent on TH, and these are strong candidates for mediators of distal patterning. Notably, we did not identify any genes or pathways that were differentially expressed in the middle of the fin where bifurcations formed. This suggests that there are not unique pathways underlying bifurcation, and that expression patterns become progressively distalized as the proximodistal axis grows.

We showed that proximally-transplanted distal portions of zebrafish fin ray tissue produced regenerates that were informed by retained memory of their original distal identity. Regenerates originating from dist-to-prox transplants retained a distal pattern of gene expression for a proximally-enriched transcript, initiated regeneration at a markedly slower pace, and regrew to a shorter length than expected for their location. Dist-to-prox transplanted rays regenerated much longer than the original size of the transplanted tissue, indicating that the proximal environment can induce considerable growth in a regenerate. Notably however, dist-to-prox regenerates grew to an ultimately much shorter length than was appropriate for their regenerating environment, and this altered length was remembered through multiple rounds of regeneration. In contrast to the regenerate length, however, we found no evidence that proximodistal patterning was remembered by dist-to-prox transplanted tissue, as these produced regenerates with segment patterning and bifurcation placement appropriate for their environmental context.

Speed of regeneration is specific to proximodistal location: proximal tissue regrows quickly while distal tissue regrows at a slower rate (Banu et al., 2022; Lee et al., 2005; Uemoto et al., 2020). Dist-to-prox transplanted rays regenerated at a slower pace during the first week of regrowth, suggesting a retained memory of distal identity. Thinner, smaller dist-to-prox rays provide fewer cells for the initial proliferation of the blastema; this smaller pool of cells may depress the initial speed of regeneration and ultimately shorten the total regenerated ray length through multiple rounds of regeneration (as in Wang et al., 2019).

Although fin rays are known to retain memory of their original length (Shibata et al., 2018), the existence of remembered identity along the proximodistal axis of the fin rays has not previously been demonstrated. Previous proximodistal transplants of blastema cells or hemi-rays did not demonstrate any retained memory of these tissues, however these transplants grafted a much smaller portion of distal tissue (Murciano et al., 2007; Shibata et al., 2018, see experimental setup diagram in Supplementary Fig. 9). There may be a threshold cell number required to specify proximodistal identity not met in these previous experiments. Our dist-to-prox transplantation relocates large numbers of numerous cell types, presumably including osteoblasts, fibroblasts, ectoderm, blood vessels, nerve tissue and other cell types—intra-ray fibroblasts are the likely mediators of positional information (Perathoner et al., 2014). Our transplant translocated sufficient types and/or numbers of cells into a proximal location to permanently alter positional memory in the regenerate.

## SUMMARY

In all, regenerating caudal fins show progressive changes in expression along the proximodistal axis, and many of these progressive changes are dependent on TH. We have shown that proximodistal gene expression patterns can be remembered autonomously by fin tissues, with dist-to-prox transplants producing regenerates with attenuated, distally-appropriate levels of *scpp7* expression. Initial rates of regenerative growth are further informed by remembered tissue identity: dist-to-prox rays begin regeneration at a slow (distally appropriate) rate. This early setback maintains the ray originating from the transplant at a shorter length than neighboring rays, and this decrease in length is remembered even through multiple rounds of regeneration.

## MATERIALS AND METHODS

### Fish rearing conditions

Zebrafish were reared at 28°C with a 14:10 light:dark cycle. Hypothyroid fish and their WT controls were *Tg(tg:nVenus-2a-nfnB)* (McMenamin et al., 2014). All other fish were WT (Tübingen line). WT fish were fed marine rotifers, *Artemia*, Gemma Micro (Skretting, Stavanger, NOR) and Adult Zebrafish Diet (Zeigler, Gardners PA, USA) 2-3 times per day. Hypothyroid fish and their WT controls were fed a diet of Spirulina flakes (Pentair, London, UK) and live *Artemia*.

### Thyroid follicle ablations

To generate hypothyroid individuals, we performed transgenic thyroid ablations (as in McMenamin et al., 2014). Briefly, to ablate the thyroid follicles of *Tg(tg:nVenus-2a-nfnB)*, 4-5dpf larvae were incubated overnight in 10 mM metronidazole (Thermo Scientific Chemicals) dissolved in 1% dimethyl sulfoxide (DMSO, Sigma) in larval water, and controls with just 1% DMSO in larval water.

### RNA Sequencing

Regenerating caudal fin tissue was collected from sibling adults (>18 standard length; SL) reared under wildtype or hypothyroid conditions during regeneration of three different regions. To minimize enrichment of genes involved in blastema formation (Li et al., 2021; Nauroy et al., 2019), we chose 4 dpa regenerates to represent proximal outgrowth. Tissue was collected at 4 dpa (proximal tissue), 7 dpa (middle tissue) or 15 dpa (distal tissue). Fish were anesthetized with tricaine (MS-222, Pentair; ∼0.02% w/v in system water), the distal-most portion of the regenerating fin (∼3 segments closest to the leading edge) was collected and immediately flash frozen in a dry ice / ethanol bath. Three or four biological replicates, each containing tissue from six individual fins, were collected at each time point and TH condition. Total RNA was extracted immediately with Zymo Quick-RNA Microprep kit R1050 (Zymo Research, Irvine CA, USA). Quality check, library preparation, and sequencing were performed by Genewiz (Cambridge, MA). Sample libraries were made with Illumina Truseq RNA Library Prep kit and sequenced on an Illumina HiSeq platform with 150bp paired-end sequence reads. Raw sequence reads were aligned to Zebrafish GRCz11 using STAR version 2.7.3 and gene counts were called with Ensembl GRCz11 gene annotation. Differential gene expression analyses were performed with Bioconductor package limma (Michaud et al., 2008). All transcriptomes were analyzed by a multidimensional scaling plot to detect overall differences in the transcriptomes. Subsequently, comparisons were made between proximal and distal regenerating regions in both WT and hypothyroid backgrounds; these were subsequently compared to identify the subset of differentially expressed genes that lost differential expression in a hypothyroid context. Genes were considered significantly expressed if they showed a log2 fold difference higher than 2 and a false discovery rate lower than 0.01.

### Microsurgeries

Transplantation was most reliable using larger adults, so all individuals used for microsurgeries were 25-40mm SL. For ray extirpation, the interray tissue on both sides of dorsal ray four (DR4) was cut (using Surgical Grade Blades #11) to separate the ray from its neighbors. The entire ray was then plucked from the peduncle, by securing the zebrafish body with General-Purpose Broad-Tipped Forceps (Fisher Scientific) while using Dumont #5 Forceps (Fine Science Tools, 1125240) to grasp the base of the ray. For dist-to-prox transplants, DR4 was extirpated from the fin, ∼2 mm of the distal tip was clipped off, and this portion was grafted back into the now-empty DR4 site (see Supplementary Fig. 10 for further detail). For prox-to-prox transplants, DR4 was extirpated and then re-inserted in its native position. Directly after transplantation, fish were maintained in a lightly anesthetized state for 30-60 minutes using ∼0.01% tricaine and 3PPM clove bud oil (Sigma-Aldrich). One day post-transplant, we assessed fins for graft success: dist-to-prox transplants grafted ∼80% of the time while prox-to-prox transplants only grafted in ∼60% of microsurgeries. After allowing 24 hours for recovery and for the graft to fuse with neighboring tissues, fish were again anesthetized with tricaine, and the entire fin (including the transplanted graft) was amputated along a single plane with a razor blade.

### RNAscope whole mount *in situ* hybridization

Regenerating fins were collected at 4 dpa (proximal tissue or dist-to-prox tissue) or 15 dpa (distal tissue) and fixed for 30 minutes in 4% PFA at room temperature. Fins were stained as described in (I. Sehring et al., 2022) with the modification that all 0.2x SSCT washes were only performed twice. We used the RNAscope Multiplex Fluorescent Reagent Kit v2 (ACD Bio-techne, 323100) to screen seven candidate probes (ACD Bio-techne: *scpp7* 1265951-C1, *rhbg* 1315181-C2, *kcnma1a* 1315191-C3, *nfil3-6* 1265961-C2, *noxo1a* 1265971-C3, *defbl1* 1265981-C4, *olfml2ba* 1315201-C4) in proximal and distal regenerating tissue. Only the *scpp7* probe was able to reliably label transcripts in our whole mount tissues. Selecting candidates with known function, gene targets were manually curated from the 45 transcripts that showed proximal or distal enrichment was dependent upon TH.

### Imaging

Zebrafish were anesthetized with tricaine and imaged on an Olympus SZX16 stereoscope with an Olympus DP74 camera or an Olympus IX83 inverted microscope with a Hamamatsu ORCA Flash 4.0 camera. Identical microscope settings (including exposure and magnification) were used for all samples within each fluorescent *in situ* experiment. Images were transformed in FIJI with the Fire LUT for visualization. For fluorescent quantifications, we used FIJI to capture mean fluorescent intensity at the distal end of dorsal ray three, dorsal ray four transplant, and dorsal ray five (DR3, DR4, and DR5).

### Analyses

All analyses were done in R 4.2.2. DR4 was used for all transplant procedures, with non-transplanted ventral ray four (VR4) serving as an internal comparison. Any damaged rays were excluded from analysis. Fin ray morphology was quantified with the StereoMorph R package (Olsen & Westneat, 2015) as described in (Harper et al., 2023). We used paired or unpaired Welch two-sample t tests or a paired repeated samples ANOVA followed by pairwise *t* tests to account for the two rays from a single fin or multiple time points assessed. Significance was marked as: p <0.05, *; p <0.01, **; p <0.001, ***.

### Pharmacological treatments

FK506 (Selleck Chemicals) was suspended in DMSO, then diluted to 200 nM FK506 and 0.02% DMSO. Controls were treated with 0.02% DMSO. ∼70% water changes were performed every other day throughout the treatment before washout. Fish recovered for seven days, then were amputated a second time with no drug treatment.

## AKNOLWEDGEMENTS

Thank you to all McMenamin Lab members past and present for fish care assistance. For data analysis support, we thank Melissa McTernan. For R assistance and dino nuggets, we thank Brian Autumn. For critical input and discussion, we thank Eric Folker, Matthew Harris, Vicki Losick and the anonymous reviewers of the manuscript.

Funding provided by NSF CAREER 1845513 and NIH R35GM146467 (S.K.M.).

## SUPPLEMENTARY FIGURES

**Supplementary Figure 1.**
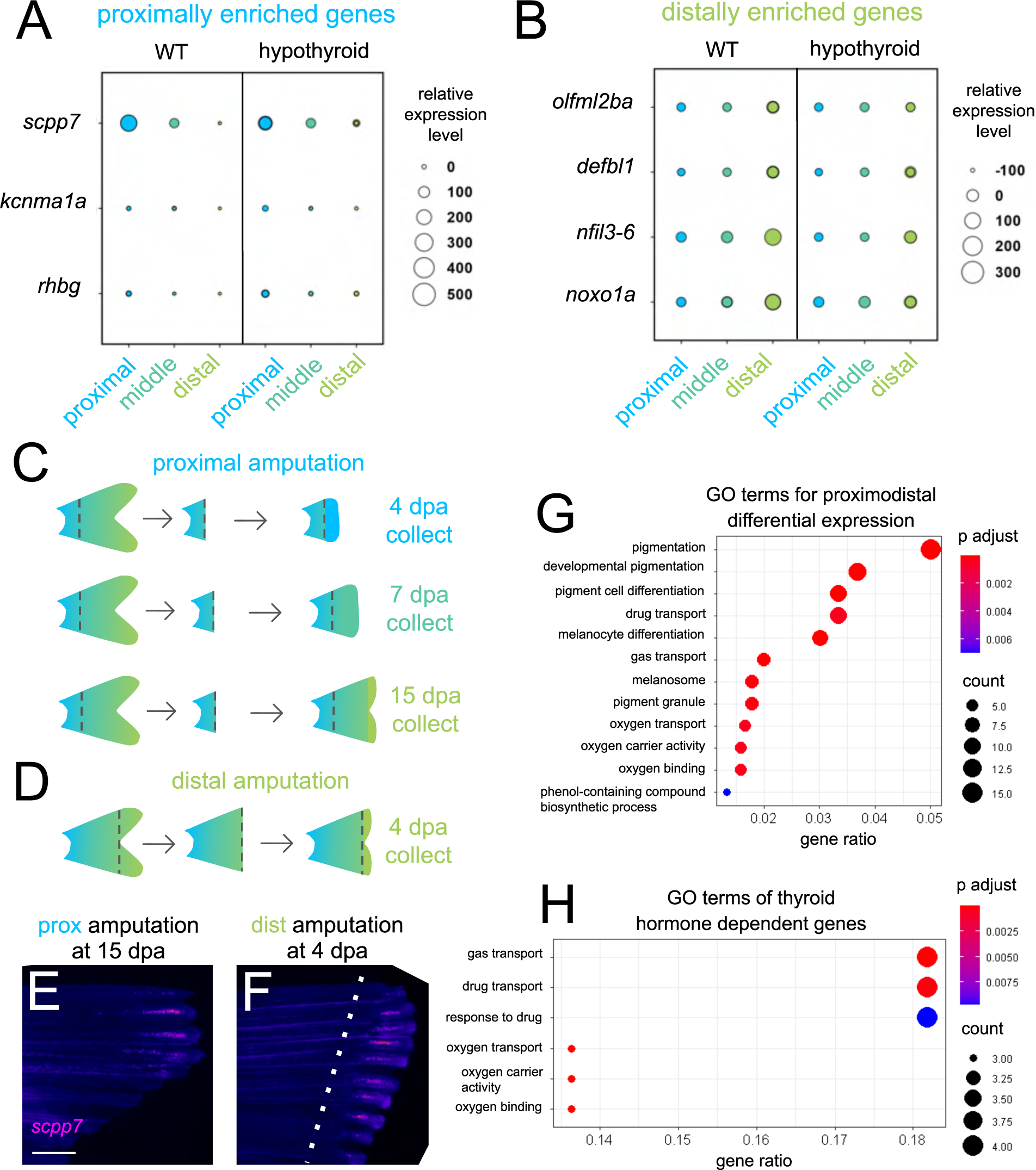
Differentially expressed gene candidates for fluorescent *in situ* hybridization. Thyroid hormone-dependent gene candidates that are either (A) proximally enriched or (B) distally enriched in WT tissues. Custom RNAscope probes were made and tested for all genes, but only the *scpp7* probe showed specific staining. (C-D) Schematic showing sample collection with (C) proximal or (D) distal amputation. (E) Proximally amputated at 15dpa or (F) distally amputated 4dpa tissue stained for *scpp7*. Amputation plane, dashed line. Warm colors indicate highest regions of expression. (G) GO enrichment of the 489 genes proximodistal differentially expressed in WT. (H) GO enrichment of the 45 genes that were thyroid hormone dependent and proximodistal differentially expressed in WT. Scale bar, 400 μm.

**Supplementary Figure 2.**
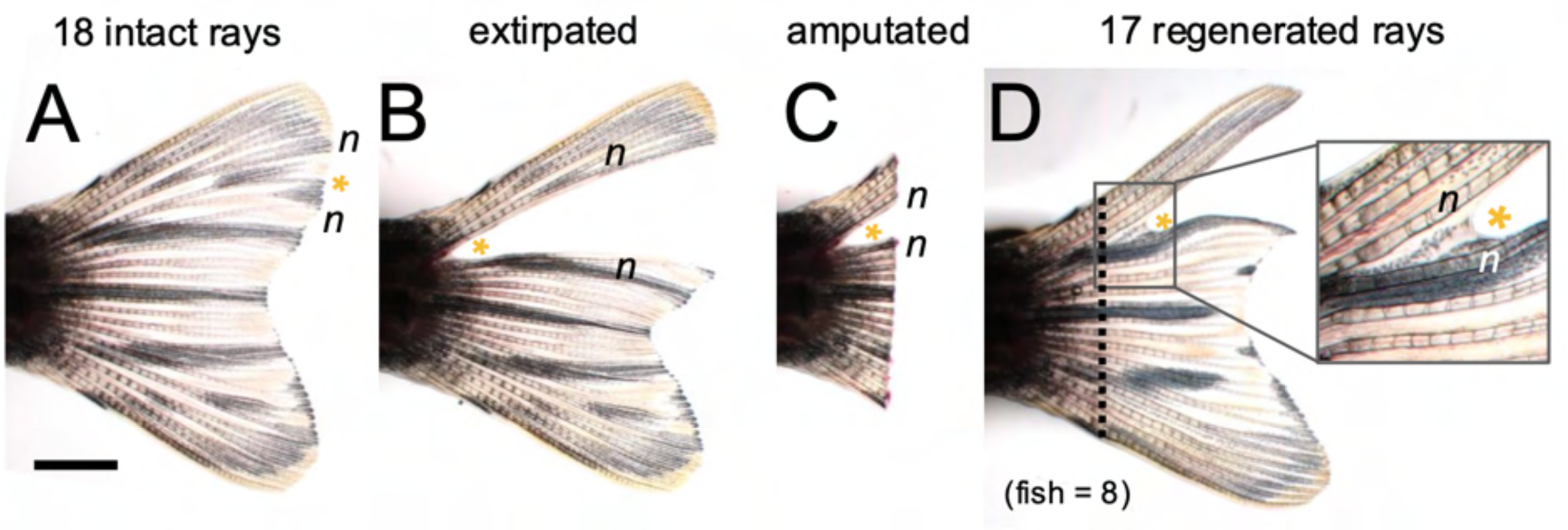
Regeneration does not originate from an extirpated ray. (A) Intact fin with 18 rays, dorsal ray 4 (D4) marked with yellow asterisk. (B) Fin one day post D4 extirpation. (C) Freshly amputated fin, one day post D4 extirpation. (D) Fin regenerates with 17 rays (one-less ray than original, intact fin). n indicates neighboring dorsal rays 3 and 5.

**Supplementary Figure 3.**
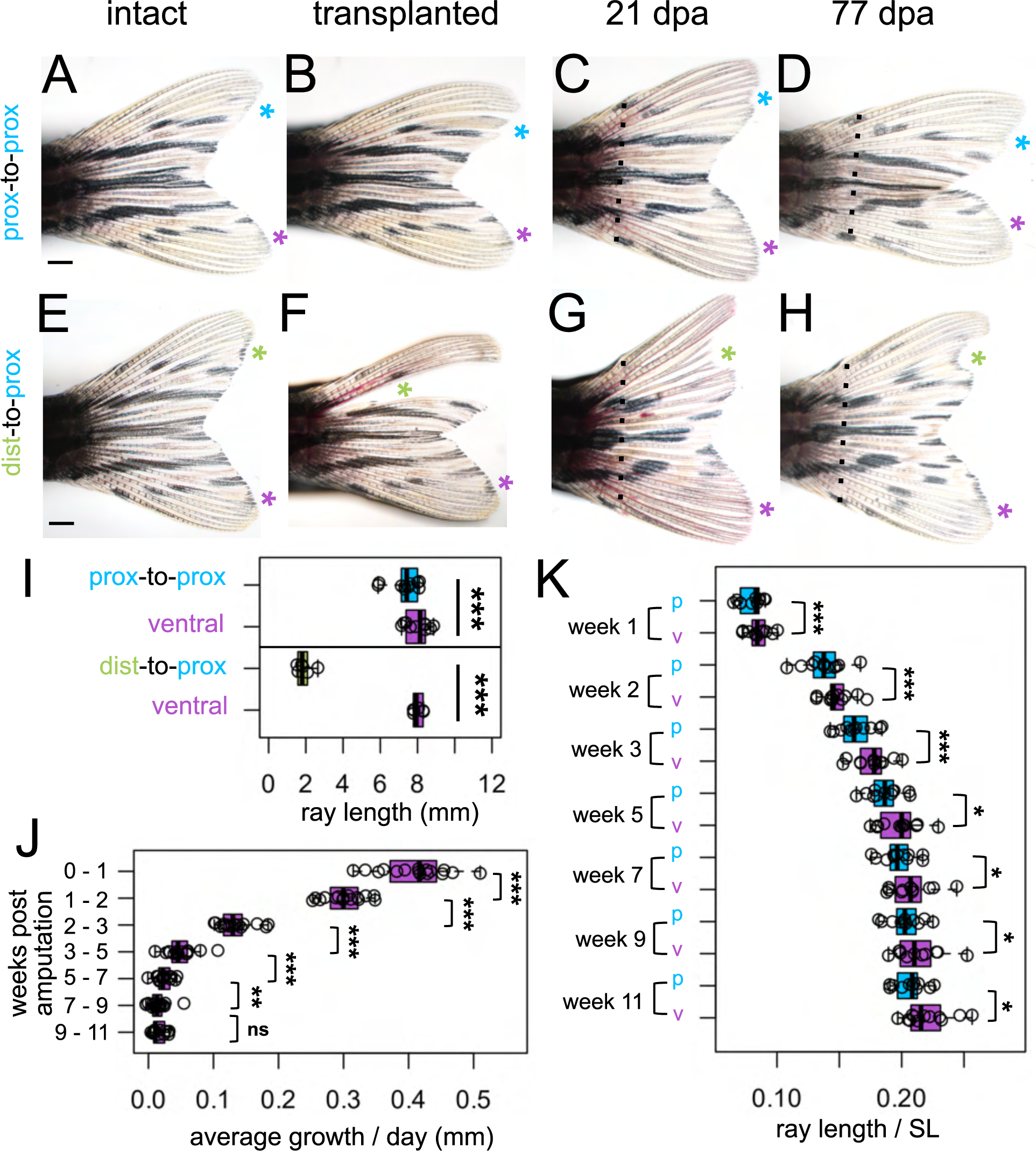
Non-transplanted rays regenerated faster than transplanted rays. Fins of (A-D) proximal-to-proximal (blue asterisk) or (E-H) distal-to-proximal (green asterisk) transplantation: (A, E) intact pre-transplantation, (B-F) one day post-transplantation, (C-G) regenerating at 21 dpa, (D, H) regenerating at 77 dpa. Ventral rays indicated with purple asterisks. Amputation plane, dashed line. (I) Length of the rays after transplantation, as measured from the peduncle. (J) Average amount of growth per day during a one/two week periods for all the ventral ray comparisons. (K) Prox-to-prox rays versus ventral ray comparisons, ray length (measured from amputation plane) divided by SL at each week. Significance determined by paired Welch two-sample t tests. Scale bar, 1 mm.

**Supplementary Figure 4.**
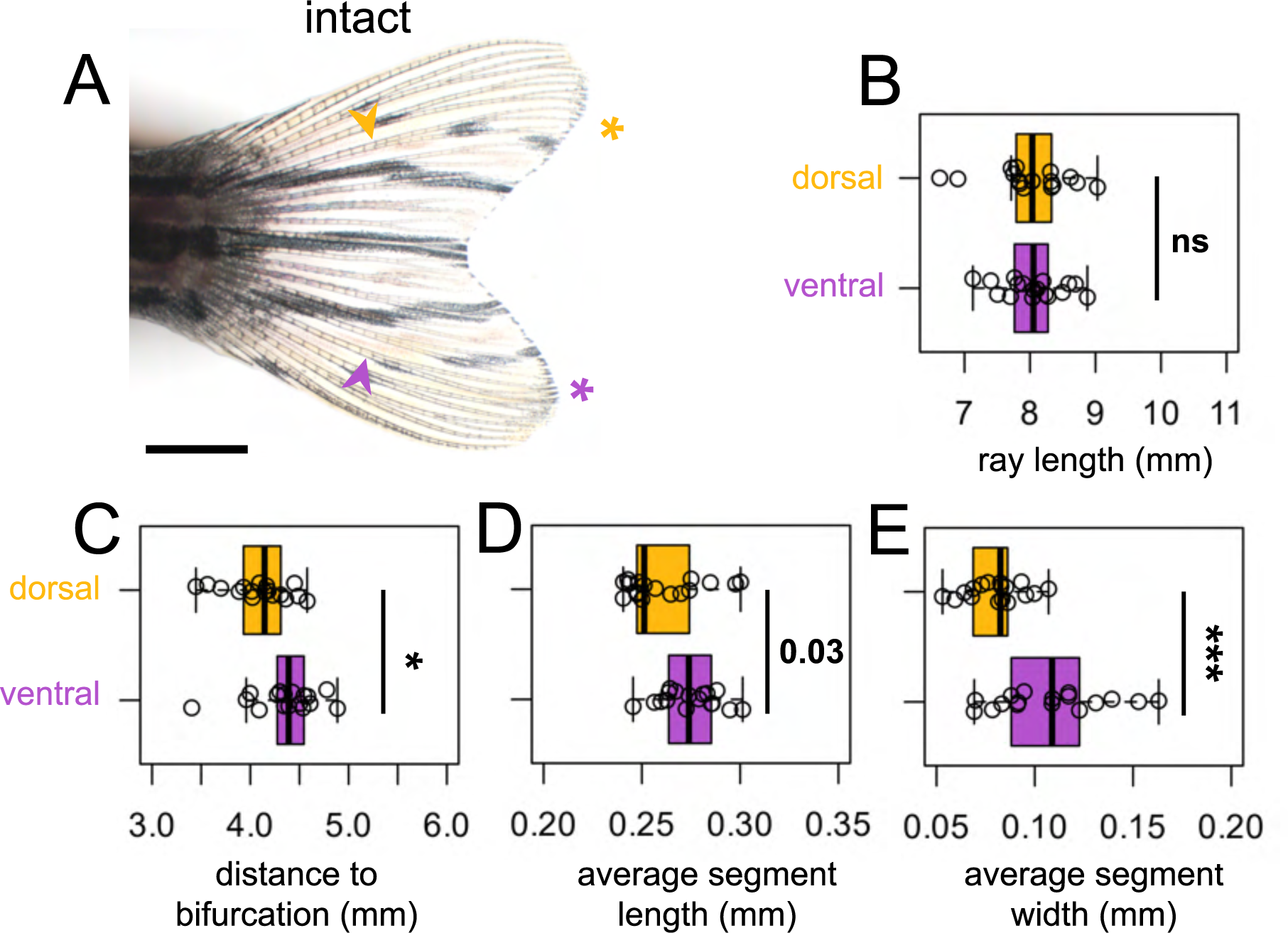
Dorsal ray patterning is unique from ventral ray patterning. (A) Intact fin. A yellow or purple asterisk indicates dorsal ray 4 or ventral ray 4, respectively. Arrowheads, primary bifurcations. Boxplots showing the (B) total length of the ray, (C) proximodistal position of the bifurcation, (D) average segment length, and (E) average segment width measured from a set distance from the peduncle. Significance determined by a paired Welch two-sample t test. Scale bar, 2 mm.

**Supplementary Figure 5.**
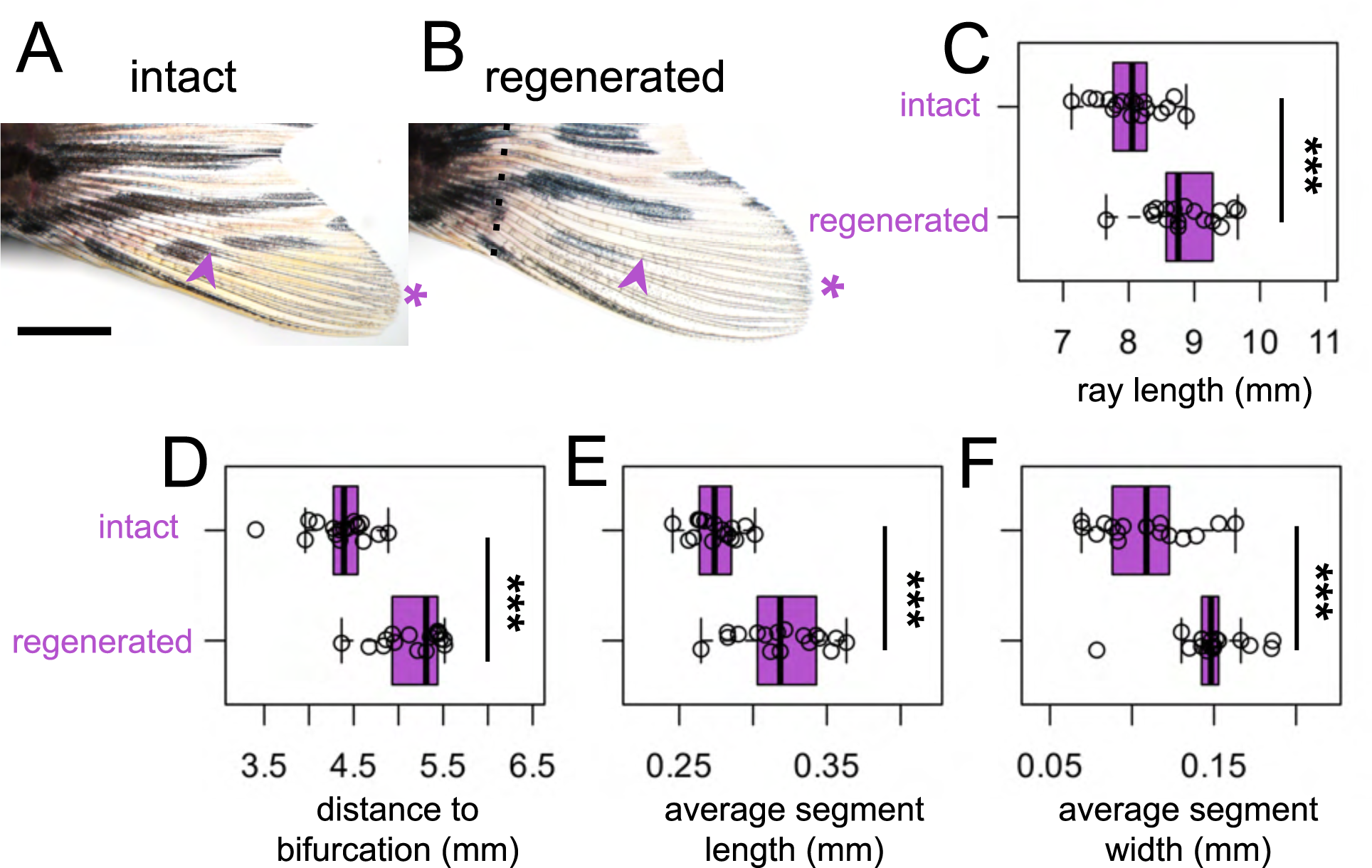
Intact and regenerated ray patterning are different. (A-B) Ventral lobe of (A) intact or (B) regenerating fin at 35dpa. Purple asterisks indicate ventral ray 4. Arrowheads, primary bifurcations. Amputation plane, dashed line. Boxplots showing the (C) total length of the ray, (D) proximodistal position of the bifurcation, (E) average segment length, and (F) average segment width measured from a set distance from the peduncle. Significance determined by a paired Welch two-sample t test. Scale bar, 2 mm.

**Supplementary Figure 6.**
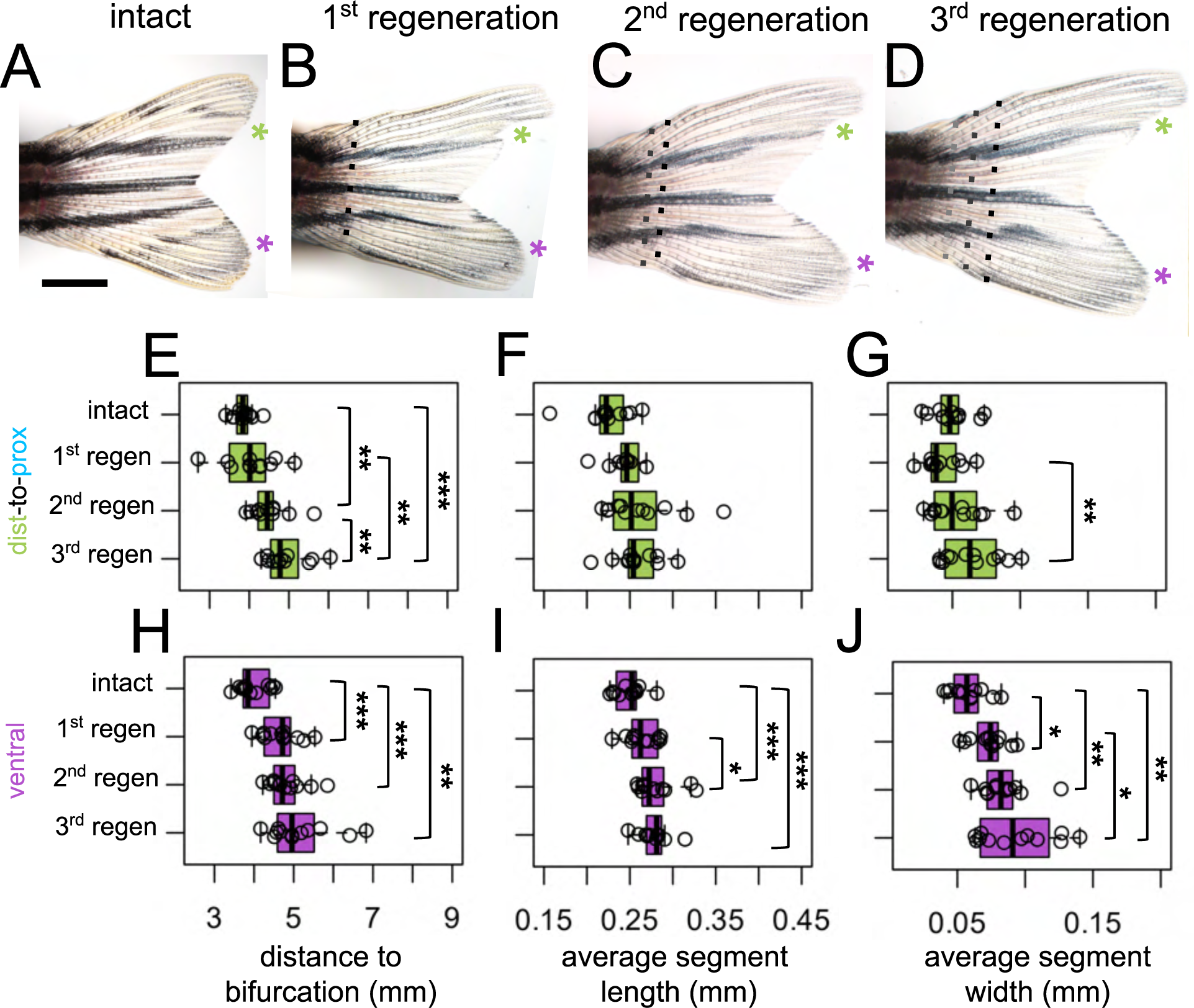
Regenerative ray patterning differs from previous regenerated morphology. (A) Intact fin. (B-D) Regenerating fin after distal-to-proximal transplantation: (B) 28 days post first amputation, (C) 28 days post second amputation, (C) 28 days post third amputation. Green or purple asterisks indicate dist-to-prox or ventral ray, respectively. Black dashed line, most recent amputation. Grey dashed lines, previous amputation planes. (E, H) Boxplots showing the proximodistal position of the bifurcation. Note that bifurcations form at increasingly distal location after each amputation, as previously described. Boxplots showing (F, I) average segment length, and (G, J) average segment width. All measurements were taken from a set distance from the peduncle. Significance determined by paired repeated samples ANOVA followed by pairwise t tests. Scale bar, 2 mm.

**Supplementary Figure 7.**
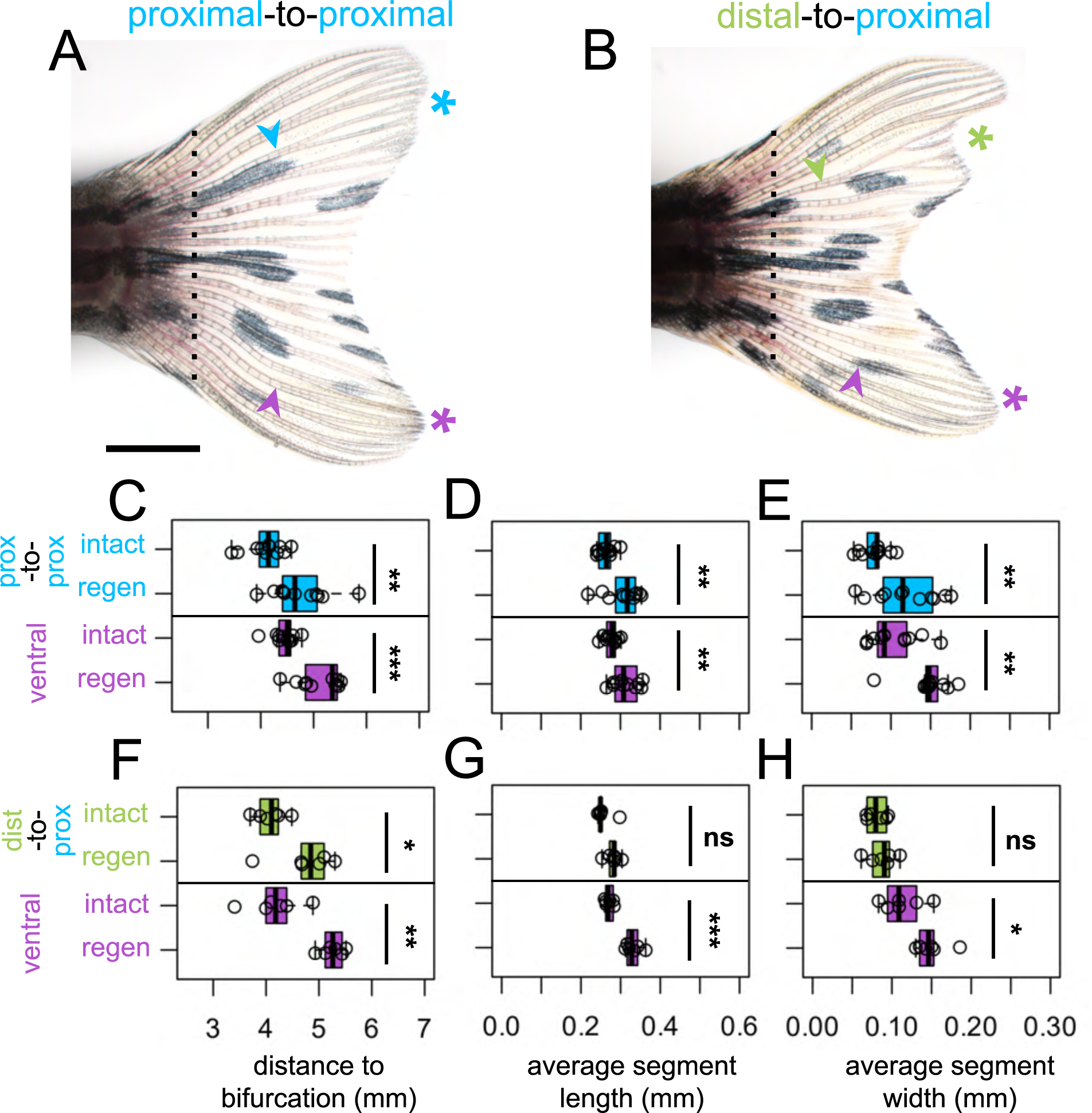
Proximodistal patterning is dependent upon the current regenerative environment. Regenerating fins at 35dpa after either (A) proximal-to-proximal (blue asterisk) or (B) distal-to-proximal (green asterisk) transplantation. Purple asterisks indicate ventral ray comparison. Amputation plane shown with dashed line. Arrowheads indicate primary bifurcations. C-H) Boxplots showing the (C, F) proximodistal position of the bifurcation, (D, G) average segment length, and (E, H) average segment width of intact or regenerated rays, measured from a set distance from the peduncle. (C-E) Prox-to-prox or dist-to-prox ray measurements are shown alongside their ventral ray comparisons. Significance determined by a paired Welch two-sample t test. Scale bar, (A-B) 2 mm.

**Supplementary Figure 8.**
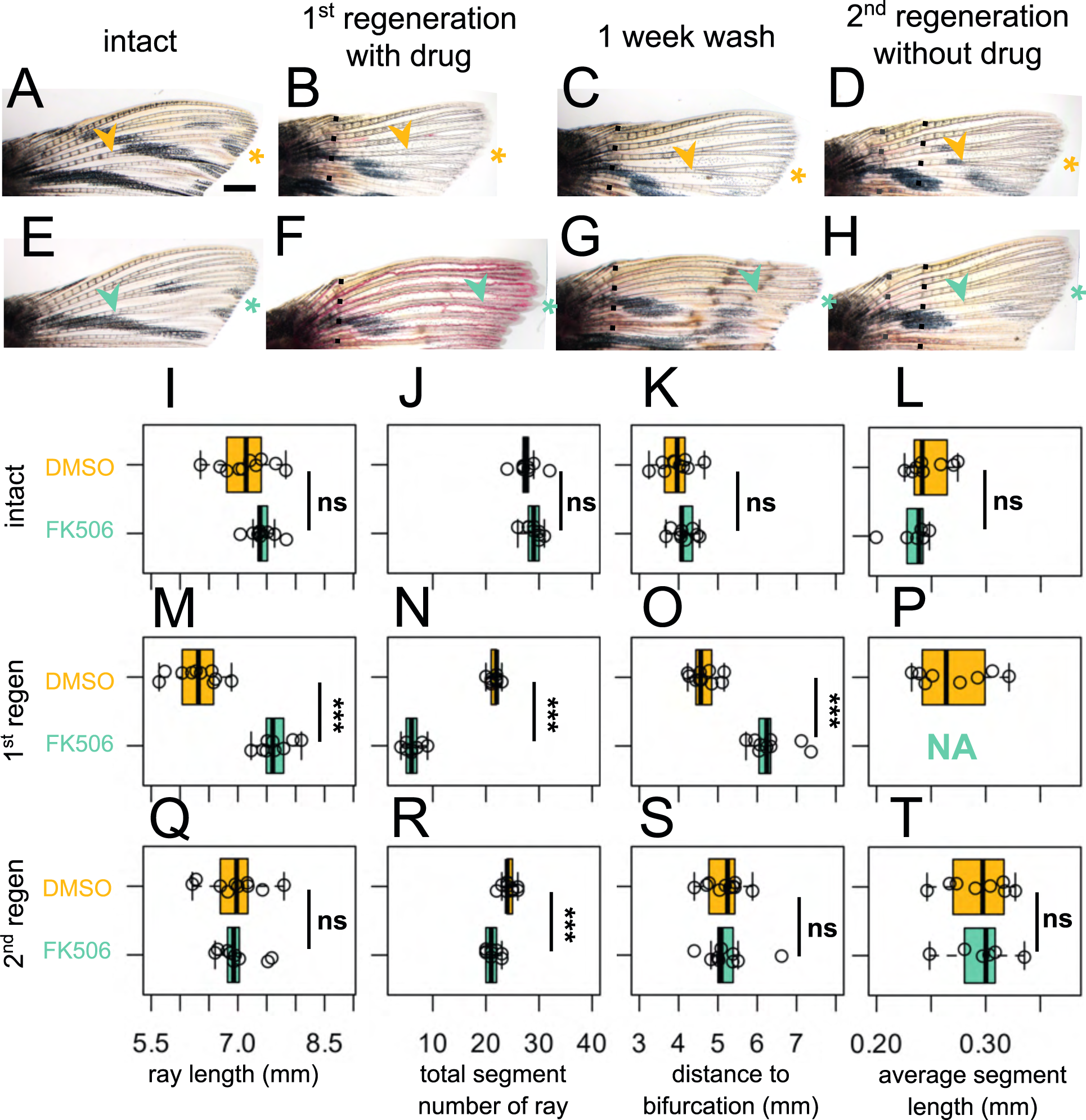
Calcineurin inhibition-induced morphologies are not remembered in subsequent regeneration cycles. (A, E) Intact dorsal lobe before treatment. (B, F) Regenerated fin after (B) DMSO (yellow asterisk) or (F) 200 nM FK506 (turquoise asterisk) treatment, 21 days post amputation. (C, G) Fins after one week wash to clear remaining drug from water. (D, H) Regenerated fin 21 days post second amputation with no treatment. Black dashed line, most recent amputation. Grey dashed lines, previous amputation plane. Boxplots showing (I, M, Q) total ray length, (J, N, R) total number of segments of the ray, (K, O, S) bifurcation position, and (L, P, T) average segment length for (I-L) intact, (M-P) first regeneration with respective drug treatment, and (Q-T) second regeneration with no drug treatment. All measurements were taken from a set distance from the peduncle. Note in (P), rays were built from only ∼5 segments, making segments lengths so long that none were contained by the standard region of interest measured. Significance determined by unpaired Welch two-sample t test. Scale bar, 1 mm.

**Supplementary Figure 9.**
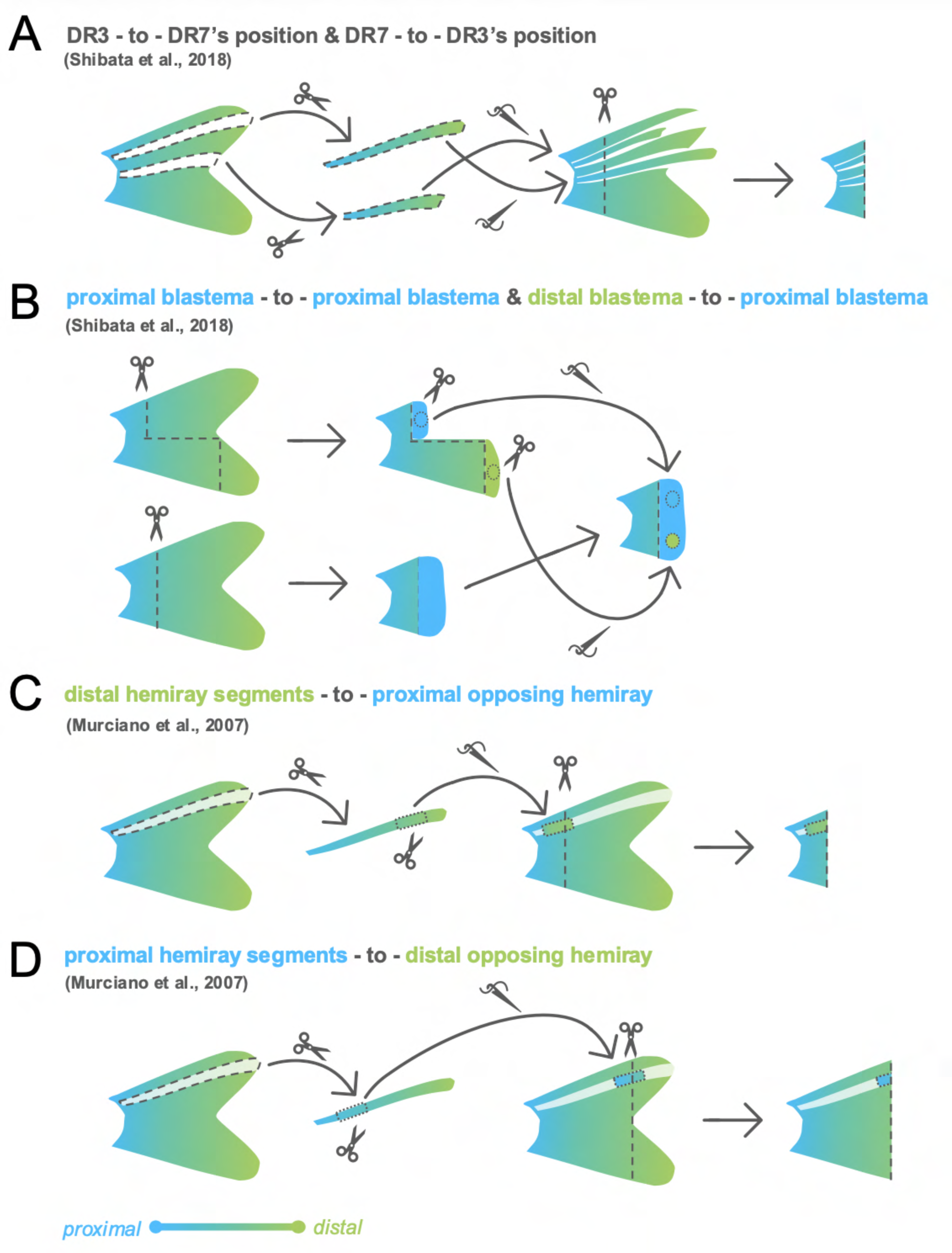
Historical transplantation experiments. (A) Shibata et al., 2018 performed full ray transplantations, moving dorsal ray 3 into dorsal ray 7 position and vise versa. After successful grafting, they amputated the entire. (B) Shibata et al., 2018 also made a proximal and distal amputation in a fin, collected blastema tissue from each region, and then transplanted these tissues into a proximally regenerating fin. (C) Murciano et al., 2007 extirpated an entire distal hemiray from the fin. A distal hemiray segment was grafted onto a proximal region to appose a proximal hemiray segment, then the entire fin was amputated through the graft. (D) Murciano et al., 2007 further extirpated a single hemiray, then grafted a proximal hemiray segment onto a distal region to appose a distal hemiray, then the entire fin was amputated through the graft.

**Supplementary Figure 10.**
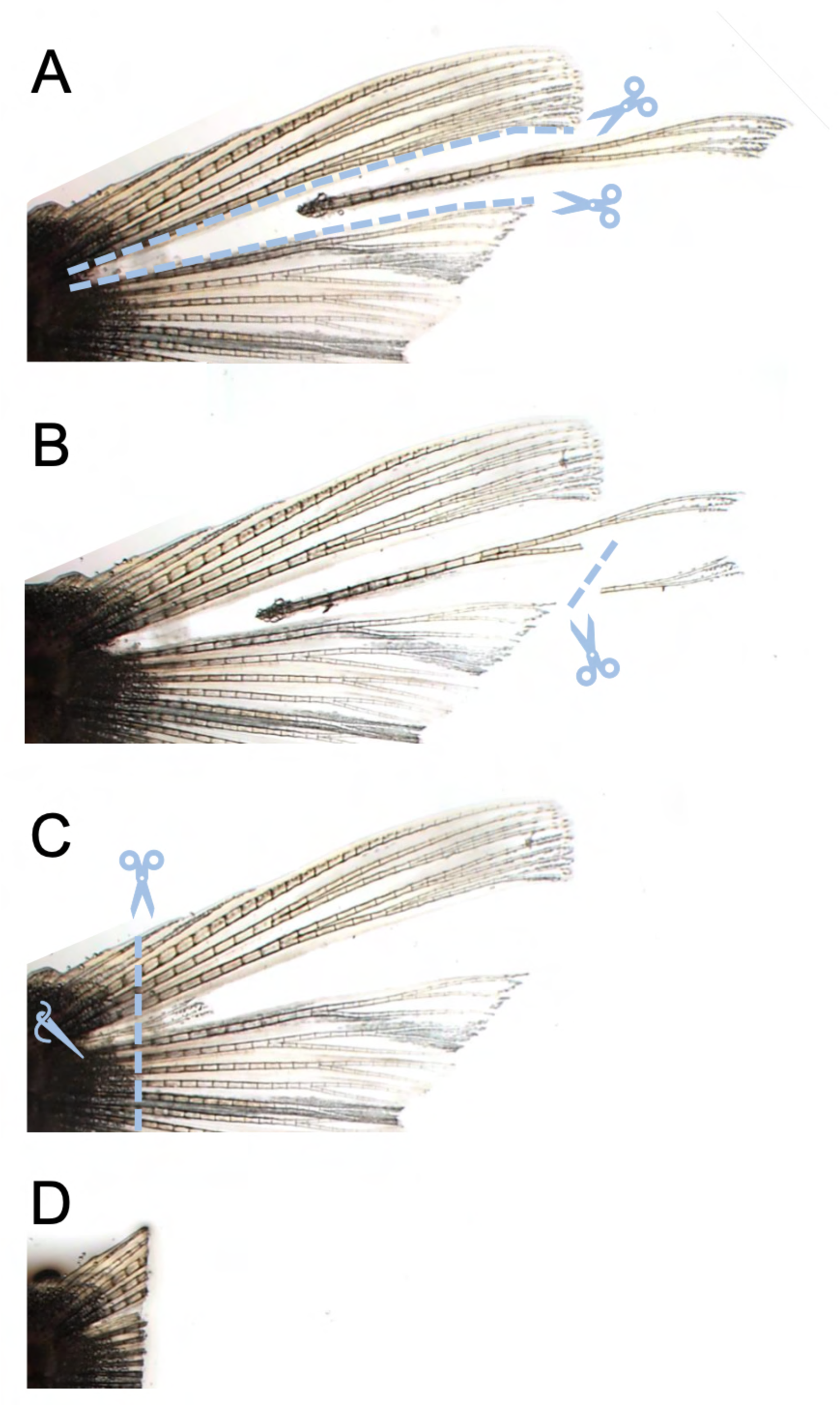
Distal-to-proximal transplantation. (A) Interray tissue is cut sliced on either side of dorsal ray 4, permitting the ray to be cleanly plucked out of the peduncle. (B) Distal ray tissue is removed from the rest of the ray. (C-D) After allowing 24hrs for the transplanted tissue to graft, the entire fin is amputated.

